# Elevation of TRPV1 expression on T cells during experimental immunosuppression

**DOI:** 10.1101/2021.03.04.434013

**Authors:** P Sanjai Kumar, Tathagata Mukherjee, Somlata Khamaru, Anukrishna Radhakrishnan, Dalai Jupiter Nanda Kishore, Saurabh Chawla, Subhransu Sekhar Sahoo, Subhasis Chattopadhyay

## Abstract

An intracellular rise in calcium (Ca^2+^) is an essential requisite underlying T cell activation and its associated pro-inflammatory cytokine production. Transient receptor potential vanilloid channel (TRPV1) is a thermo-sensitive, polymodal gated and permeable to cations such as Ca^2+^. It has been reported that TRPV1 expression increases during T cell activation. However, the possible involvement of TRPV1 during immunosuppression of T cells has not been studied yet. Here, we investigated the possible role of TRPV1 in FK506 or B16F10-culture supernatant (B16F10-CS) driven experimental immunosuppression in T cells. Intriguingly, it was found that TRPV1 expression is further elevated during immunosuppression compared to ConA or TCR activated T cells. Similarly, in B16F10 tumor-bearing mice, the TRPV1 expression was upregulated in T cells as compared to control mice, *in vivo*. Moreover, we observed an immediate rise in intracellular Ca^2+^ levels in FK506 and B16F10-CS treated T cells as compared to ConA or TCR treated T cells. Likewise, in B16F10 tumor-bearing mice, the basal intracellular calcium level was upregulated in T cells as compared to control mice, *in vivo*. To further investigate the possible mechanism of such rise in intracellular Ca^2+^ levels, TRPV1 specific functional inhibitor, 5 -iodoresiniferatoxin (5 -IRTX) was used in calcium influx studies. It was observed that the total intracellular Ca^2+^ levels decreased significantly in presence of 5 -IRTX for either the FK506 or B16F10-CS as well as with ConA or TCR stimulated T cells, indicating the functional role of TRPV1 channels in FK506 or B16F10-CS mediated increase in intracellular Ca^2+^ levels. The current findings highlight an essential role of the TRPV1 channel in upregulating intracellular calcium levels during both immune-activation and immunosuppression. This study might also have broad implications in the context of other immune-suppressive diseases as well.

## 1. Introduction

Transient receptor potential channels (TRP) belong to a superfamily of ion channels that are thermo-sensitive, polymodal gated and permeable to cations (Gees et al., 2010; Nilius, 2007; Samanta, Hughes, & Moiseenkova-Bell, 2018). Transient receptor potential vanilloid channels (TRPV1) are reported to have a regulatory role in various T cell processes including differentiation, proliferation and activation (Amantini et al., 2017; Majhi et al., 2015). Activation of TRPV1, via ligand binding, significantly increases TCR mediated Ca^2+^ influx and downstream signaling pathways (Majhi *et al*., 2015; Bujak *et al*., 2019). TRPV1 is distributed widely in immune cells and are involved in various inflammatory conditions such as in inflammatory bowel disease (IBD), cutaneous neurogenic inflammation, chronic obstructive pulmonary disease (COPD), allergic asthma, brain inflammation, arthritis, auto-immune diseases, hypersensitivity and colitis (Bassi et al., 2019; Silva et al., 2018) and various viral infections as well (Bassi et al., 2019; Kumar et al., 2020; Omar et al., 2017; Omari et al., 2017; Silva et al., 2018).

Free Ca^2+^ ions serve as a second messenger of cells including different immune cells such as T and B lymphocytes and macrophages regulating various physiological processes like differentiation, gene transcription and effector function (Oh-hora & Rao, 2008; Vig & Kinet, 2009). Receptor-mediated activation of different immune cells has been reported to upregulate intracellular Ca^2+^ levels (Vig & Kinet, 2009). Abnormality in Ca^2+^ signaling or unavailability in immune cells may lead to various immunological disorders like severe combined immunodeficiency (SCID) and Wiskott–Aldrich syndrome (WAS) (Feske, 2007) or hamper proper activation of naive T cell and effector function (Birx et al., 1984; Nakabayashi et al., 1992; Watman, et al., 1988) respectively. Different intracellular signaling members of T cells like PKC, NFAT, NF-kB, calmodulin-dependent kinase, JNK require Ca^2+^ for their function (Ando et al., 2014; Komada et al., 1996; Vig & Kinet, 2009).

Concanavalin A (ConA), a plant lectin (carbohydrate-binding protein), extracted from Jack Bean (*Canavalia ensiformis*), acts as a mitogen for T cell (Mackler et al., 1972) and can increase intracellular Ca^2+^ concentration required for IL-2R expression and IL-2 production (Komada et al., 1996). ConA binds to the mannose moieties of glycoproteins and glycolipids, including the T cell receptor (TCR) and is responsible for nonspecific T cell proliferation (Ando et al., 2014). On the other hand, T cell activation via TCR stimulation with anti-CD3 and anti-CD28 is more specific proliferation which also results in the rise of intracellular Ca^2+^ levels (Pang et al., 2012). Collectively, ConA or TCR induced increase of intracellular Ca^2+^ involves PLC/InsP3, which releases Ca^2+^ from intracellular compartments like the endoplasmic reticulum and then opens plasma membrane residing Ca^2+^ channels like calcium release-activated calcium channel (CRAC) and different TRP channels (Pang *et al*., 2012; Majhi *et al*., 2015).

FK506 (tacrolimus), a clinical immunosuppressive agent, binds calcineurin and impede pro-inflammatory cytokine production by T cells (Almawi et al., 2001; Annett et al., 2020; Sakuma et al., 2000). FK506 binds to FK506 binding protein (FKBP) and forms complex FK506-FKBP. FK506 binds to FK506 binding protein (FKBP) and forms FK506-FKBP complex (Bultynck et al., 2000). FKBPs are reported as important modulators of intracellular Ca^2+^. FKBPs regulate intracellular ryanodine and IP3 receptor Ca^2+^ release channels (Bultynck et al., 2000; Cameron et al., 1995; MacMillan, 2013). Recently, it has been reported that FK506 induces Ca^2+^ influx via TRPA1 channels (Bultynck et al., 2000; Kita et al., 2019).

B16F10, a mouse melanoma cell line, has been widely used as a transplantable murine melanoma model (Burghoff et al., 2014; Chen et al., 2019; Langer et al., 2006; Overwijk & Restifo, 2001). Additionally, it has also been reported to secrete various soluble factors from B16F10 derived tumors, suppressing lymphocyte activation *in vitro* (Sun et al., 2015, 2013, 2011). B16F10 derived soluble factors, such as interleukin-10 (IL-10), transforming growth factor-beta (TGF-β), vascular endothelial growth factor (VEGF), play an important role in immunosuppression. These soluble factors are known to develop an immunosuppressive network spanning from the primary tumor site to secondary lymphoid organs and peripheral vessels (Yang & Carbone, 2004; Zou, 2005). Tumor microenvironment milieu enriched with these soluble factors induces immune cells to become immunosuppressive and hinders T cell activation associated with antitumor responses (Kusmartsev & Gabrilovich, 2002; Kusmartsev et al., 2004). Additionally, tumor-released extracellular vesicles from the supernatant of malignant effusions or tumor cells such as B16F10 and ascites of cancer patients are also reported to induce immunosuppression. Immune cells such as T cells, B cells, neutrophils and macrophages can take up these tumor-released extracellular vesicles and induce an immunosuppressive environment (Chen et al., 2019; Gao et al., 2018; Wen et al., 2018; Zhou et al., 2016).

The functional role of TRPV1 in cell-mediated immunity (CMI) and its role towards an increase in intracellular Ca^2+^ has been reported (Bertin *et al*., 2014; Majhi *et al*., 2015; Kumar *et. al*., 2020). Moreover, a number of immunosuppressive agents, such as rapamycin, tacrolimus and cyclosporin has also been reported to induce intracellular rise in Ca^2+^ (Bielefeldt, Sharma, Whiteis, Yedidag, & Abboud, 1997; Bultynck et al., 2000; Cameron, Steiner, Roskams, et al., 1995; Cameron, Steiner, Sabatini, et al., 1995; Kanoh et al., 1999; D. MacMillan & McCarron, 2009; Van Acker et al., 2004). Based on these previous findings, we hypothesized that TRPV1 may also be contributing towards the intracellular rise in Ca^2+^ levels, subsequent immunosuppressive downstream signaling and cellular responses. Therefore, in the present study, the possible role of TRPV1 during experimental immunosuppression was explored. Moreover, FK506 or B16F10-CS driven regulation in T cell activation, inflammatory cytokines response and subsequent Ca^2+^ influx was investigated. The current study might have importance in understanding the TRPV1 driven FK506 or B16F10-CS induced immunosuppression in T cells, and it might have implications in various immunosuppressive diseases as well.

## 2. Materials and Methods

### 2.1 Mice

6 to 8 weeks old both male and female C57BL/6 mice from the National Institute of Science Education and Research (NISER), Bhubaneswar were used for experimentation. All protocols were approved by the Institutional Animal Ethics Committee (IAEC), NISER following Committee for the Purpose of Control and Supervision of Experiments on Animals, CPCSEA guidelines (IAEC protocol no. AH-112).

### 2.2 Antibodies, reagents and drugs

FK506 (Cat. no. F4679-5MG), Concanavalin A (ConA) (Cat. no. C0412-5MG), 5 -IRTX (Cat. no. I9281-1MG) were purchased from Sigma Aldrich (St. Louis, MO, USA); anti-mouse CD25 (Cat. no. 553866), OptEIA kits for IL-2 (Cat. no.555148), IFN-γ (Cat. no.555138), TNF (Cat. no. 558534) for sandwich ELISA were from BD Biosciences, SJ, CA, USA.; anti-mouse CD69 (Cat. no. 35-0691-U100), anti-mouse CD90.2 (Cat. no. 20-0903-U100) from Tonbo Biosciences, San Diego, CA, USA; anti-mouse TRPV1 (Cat. no. ACC-029) from Alomone Laboratories (Jerusalem, Israel); anti-mouse CD3 (Cat. no. 40-0032-U500) and anti-mouse CD28 (Cat. no. 40-0281-U500) [functional assay grade] were procured from Tonbo Biosciences, San Diego, CA, USA. Secondary anti-rabbit Alexa Fluor 488 and T cell isolation kit (Dynabeads™ Untouched™ Mouse T Cells Kit, Cat. no. 11413D) were procured from Invitrogen, Carlsbad, CA USA. RPMI-1640 cell culture medium and FBS were purchased from PAN Biotech, Aiden Bach, Germany; DMEM cell culture medium, 10X RBC lysis buffer, 10X PBS, L-glutamine, penicillin, streptomycin was from Himedia Laboratories, Mumbai, Maharashtra, India.

### 2.3 T cell isolation, purification and cell culture

Spleens were collected from C57BL/6 mice and splenocytes were isolated as mentioned previously (Sahoo *et al*., 2018). In brief, spleens were mechanically dispersed with a syringe plunger (Dispovan) and passed through a 70 uM cell strainer (SPL Life Sciences, Korea). After centrifugation, RBCs were lysed using RBC lysis buffer, washed with 1X PBS, centrifuged and resuspended in RPMI-1640 medium supplemented with 10% FBS, 2 mM L-glutamine, 100 U/mL penicillin, and 0.1 mg/mL streptomycin. T cells were then purified using Mouse Untouched T cell isolation kit as per manufacturer protocol. Briefly, splenocytes were incubated with biotinylated antibodies for 20 mins following resuspension in isolation buffer (2% FBS, 2 mM EDTA in 1X PBS). Cells were then washed with excess isolation buffer, centrifuged and incubated for 15 minutes with streptavidin-conjugated magnetic beads and then placed in a magnet for 2 minutes. Purity of T cells was ≥95% measured using flow cytometry (BD FACSCalibur™ flow cytometer, BD Biosciences).

B16F10 (ATCC^®^ CRL-6475™) cells were cultured in DMEM supplemented with 10% FBS, 2 mM L-glutamine, 100 U/mL penicillin, and 0.1 mg/mL streptomycin and maintained as per ATCC protocol. For B16F10 culture supernatant (B16F10-CS) collection, 1×10^6^ B16F10 cells were seeded in a T-75 cell culture flask (SPL Life Sciences, Korea). When the confluency reaches around 60-70%, culture medium was replaced with fresh medium and allowed to grow for 24 hours. Culture medium was then collected, centrifuged, filtered through 0.22 µM filter and stored at −80°C till further use. Purified T cells [1.0×10^6^ cells per well in a 48 well cell culture plate (SPL Life Sciences, Korea), 1 mL total volume] were then stimulated with either ConA (5 µg/mL) or anti-CD3 (plate-bound) and anti-CD28 (soluble) (2 µg/mL each) following pretreatment with FK506 (1 hour) or B16F10-CS inside a humidified incubator with 5% CO2 and 37°C temperature. After 36 hours, cells were harvested, stained and analyzed via flow cytometry. Cell culture supernatants were collected and stored at −20°C for ELISA.

### 2.4 B16F10 subcutaneous injection

B16F10 cells in the logarithmic growth phase (≤50% confluency) were harvested and suspended in ice-cold HBSS buffer. After counting, 2 × 10^5^ cells/mouse were injected in the right flank of wild type C57BL/6 mice. From 14^th^ day, tumor growth was periodically monitored in an interval of every second day (Overwijk & Restifo, 2001). On 21^st^ day, the mice were sacrificed and spleen was harvested for further processing.

### 2.5 Cell Viability Assay

Purified mouse T cells were incubated with different doses of B16F10-CS (5 to 40% of final volume), 5 -IRTX (2.5 to 20µM) and FK506 (1.25 to 20µg/mL) for 36 hours. Next, cells were harvested and washed with 1X PBS, followed by incubation with 7-AAD for 15 minutes at RT. Cells were then acquired in BD LSRFortessa™ (BD Biosciences) and analyzed via FlowJo V10.7.1. 7-AAD negative cells were considered as live cells (Nayak et al., 2017; Sahoo et al., 2018).

### 2.6 Flow Cytometry

Flow cytometric staining of T cell was performed as per the method reported previously (Majhi *et al*., 2015; Sahoo *et al*., 2018, 2019). Cells were harvested, washed with 1X PBS, resuspended in FACS staining buffer (1% BSA, 0.01% NaN3 in 1X PBS) and incubated with fluorochrome-conjugated antibodies for 30 minutes on ice in dark and then washed twice with FACS staining buffer. For TRPVI staining only, secondary fluorochrome-conjugated AF488 was added and incubated for 30 minutes, followed by washing with FACS buffer. Cells were then fixed and acquired using the BD FACSCalibur™ flow cytometer or BD LSRFortessa™ (BD Biosciences) and analyzed via Flowjo V10.7.1.

### 2.7 Enzyme-linked Immunosorbent Assay (ELISA)

Sandwich ELISA was performed to quantitate the cytokine levels in cell culture supernatants. ELISA for IL-2, IFN-γ and TNF were performed using BD OptEIA ELISA kits as per manufacturer’s protocol (Kumar et al., 2020; Sahoo et al., 2019). The cytokine concentration in supernatants was estimated by comparing the corresponding standard curve using different concentrations of the recombinant cytokines in pg/ml.

### 2.8 Ca^2+^ influx

Ca^2+^ influx in purified splenic T cells was performed as reported elsewhere (Kume & Tsukimoto, 2019; Majhi et al., 2015; Sahoo et al., 2019) Here in brief, purified splenic T cells were loaded with Ca^2+^ sensitive dye (Fluo-4 AM, 2 μM) for 60 minutes at 37°C. Next, T cells were washed twice with 1X PBS and placed inside the incubator for another 30 minutes for de-esterification. The cell suspension was added to RIA vials and then treated as per the experimental conditions and acquired by Flow cytometer. Such values were plotted for fluorescence signal relative to starting signal as (F/F0).

To estimate the intracellular Ca^2+^ contributed by the TRPV1 channel, we performed a minor modification to the Fluo-4 AM staining protocol. The minor modification has been detailed below. First, the T cells were loaded with Ca^2+^ sensitive dye (Fluo-4 AM, 2 μM) along with FK506 (5µg/mL), B16F10-CS (20%), ConA (5µg/mL) or TCR (2µg/mL of anti-CD3 and anti-CD28) for 60 minutes at 37 °C, as per the experimental conditions (instead of adding the above reagents, during acquiring). Additionally, another similar set-up of T cells were pre-treated with 5 -IRTX (10µM) for 15 minutes before addition of the FK506, B16F10-CS, ConA or TCR (same amounts as mentioned above) for 60 minutes at 37 °C, as per the experimental conditions. Next, T cells were washed twice with 1X PBS and placed inside an incubator for another 30 minutes for de-esterification. The cell suspension was added to the RIA vials for the measurement of accumulated intracellular Ca^2+^ levels. The cells were acquired in flow cytometer. The rationale behind the idea was to test whether the treatment with these reagents (FK506, B16F10-CS, ConA or TCR) lead to rise in intracellular Ca^2+^ over time whereas T cells pre-treated with 5 -IRTX (Kumar et al., 2020; Majhi et al., 2015) may regulate intracellular Ca^2+^ in T cells.

### 2.9 Statistical Analysis

Statistical analysis was performed using the GraphPad Prism 7.0 software (GraphPad Software Inc., San Diego, CA, USA). Data are represented as Mean ± SEM. The comparison between the groups was performed by one-way ANOVA or student s t-test with the sidak posthoc test. Data presented are representative of at least three independent experiments. Statistical significance is represented by asterisks (*) for p-value(s) and is marked correspondingly in the figures (\**p*< 0.05, \*\**p*<0.01, \*\*\**p*<0.001)

## 3. Results

### 3.1 Upregulation of TRPV1 during immune-activation and immunosuppression

A number of TRP channels are known to be expressed on T cells including TRPV1, TRPV4, TRPA1, TRPM8 (Acharya et al., 2020; Bertin et al., 2014; Majhi et al., 2015; Sahoo et al., 2019). To determine the expression of TRPV1 channels in mouse splenic purified resting T cells, we performed Flow cytometry, using specific antibodies against TRPV1. It was observed that the TRPV1 percent positive cells were 17.86 ± 1.34 as compared to isotype control (0.02 ± 0.01) (Supplementary Figure 1, A). Further, the specificity of the TRPV1 antibody in T cells was tested by using control blocking peptide antigen. For that, T cells were stained with anti-TRPV1 antibody in the presence or absence of the blocking peptide. It was observed that the percentage of positive cells for TRPV1 was markedly reduced in a dose-dependent manner from 17.86 ± 1.34 to 1.46 ± 0.11 in the presence of 1X control blocking peptide antigen and further reduced to 0.74 ± 0.09 in the presence of 3x control blocking peptide antigen. These results indicate that TRPV1 is expressed on T cells and the anti-TRPV1 antibody is highly specific towards TRPV1 expressed on T cells (Supplementary Figure 1, A).

7-aminoactinomycin D (7-AAD) is a fluorescent intercalator that undergoes a spectral shift upon association with DNA. It is one of the widely used cell viability dye (Nayak et al., 2017; Sahoo et al., 2018). In order to determine the cellular cytotoxicity of 5 -IRTX, FK506 and B16F10-CS in purified mouse T cells, we performed 7-AAD staining. It was observed that upto 96% of the cells were 7-AAD negative for 5 -IRTX in a concentration range of 2.5 µM to 20 µM. Similarly, approximately upto 96% of the cells were 7-AAD negative for FK506 in a concentration range of 1.25 µg/ml to 20 µg/ml. Likewise, approximately upto 95% of the cells were 7-AAD negative for B16F10-CS in a range of 5% to 40% w.r.t. total media volume (Figure 1, A). In all the experiments, heat-killed purified T cells were used as a positive control. For further experiments, we have chosen concentration with upto 95% viability. Accordingly, we have chosen 10 µM 5 -IRTX, 5 µg/ml FK506 and 20 % of B16F10-CS.

**FIGURE 1:**
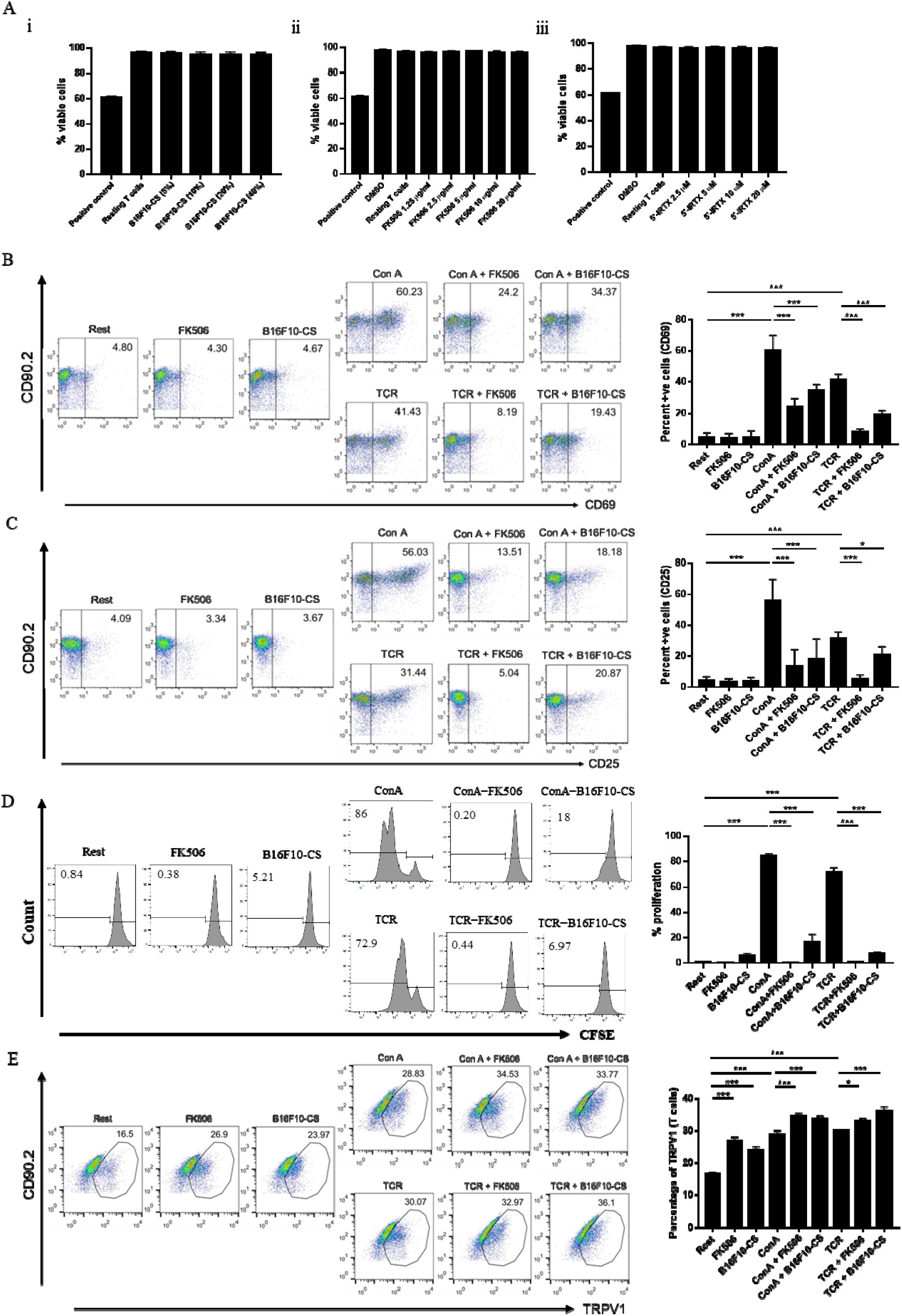
Upregulation of TRPV1 during immune-activation and immunosuppression. (A) T cells viability upon treatment with different concentrations of either (i) FK506, (ii) B16F10-CS or (iii) 5 -IRTX, as assessed by 7-AAD staining. FC dot-plot depicting T cell activation markers (B) CD69 and (C) CD25 along with its corresponding bar diagram. (D) Histogram representing T cell proliferation as determined by CFSE staining along with its corresponding bar diagram. (E) FC dot-plot showing TRPV1 expression in T cells along with its corresponding bar diagram. Representative data of three independent experiments are shown. *P* < 0.05 was considered as statistically significant difference between the groups (ns, non-significant; * *p* < 0.05; ** *p* < 0.01; *** *p* < 0.001).

TRPV1 has been attributed in functional activation and regulation of T cells (Bertin et al., 2014; Majhi et al., 2015; Omari et al., 2017). Recently, TRPV1 has also been reported to play an active role in various inflammatory and autoimmune diseases (Bassi et al., 2019). To determine the status of T cell activation during immunosuppressive treatment with either FK506 or B16F10-CS, T cell activation markers including CD69 and CD25 were analyzed via Flow cytometry. For immunosuppression, T cells were treated with either FK506 or B16F10-CS for 1 hour prior to the addition of either ConA or TCR. It was observed that in T cells treated with FK506 or B16F10 in presence of ConA, CD69 decreased significantly to 24.2 ± 5.36 and 34.37 ± 3.85 respectively as compared to control ConA activated cells (60.23 ± 9.64). Similarly, treatment with FK506 or B16F10 in presence of TCR stimulations, CD69 decreased significantly to 8.19 ± 1.93 and 19.43 ± 2.45 respectively as compared to control TCR activated cells (41.43 ± 3.37) (Figure 1, B). These results indicate that CD69 significantly decreases in FK506 or B16F10-CS treated T cells stimulated with ConA or TCR.

Correspondingly, in FK506 or B16F10-CS treated T cells, stimulated with ConA, CD25 decreased significantly to 13.51 ± 10.65 and 18.18 ± 12.94 respectively as compared to control ConA activated cells (56.03 ± 13.51). Similarly, FK506 or B16F10-CS treated T cells stimulated with TCR, CD25 decreased significantly to 5.03 ± 2.92 and 20.87 ± 5.31 respectively as compared to control TCR activated cells (31.44 ± 4.21) (Figure 1, C). These results indicate that CD25 significantly decreases in FK506 or B16F10-CS treated T cells stimulated with ConA or TCR.

T cell activation is accompanied by proliferation of T cells. To ascertain whether immunosuppressed T cell’s stimulation via either ConA or TCR leads to reduced proliferation, we performed T cell proliferation assay via CFSE staining. It has been observed that 84.53 ± 1.33 % of cells proliferated in presence of ConA. In T cells pre-treated with FK506 or B16F10-CS and later stimulated with ConA, we found that 0.32 ± 0.18% of cells and 16.5 ± 6.09% of cells proliferated, respectively. Similarly, it has been observed that 71.1 ± 3.74% of cells proliferated in presence of TCR. In T cells pre-treated with FK506 or B16F10-CS and later stimulated with TCR, we found that 0.62 ± 0.22% of cells and 7.62 ± 0.83% of cells proliferated, respectively (Figure 1, D). These results indicate that immunosuppression via either FK506 or B16F10-CS attenuates T cell proliferation.

Subsequently, the expression of TRPV1 was assessed in immunosuppressed T cells. Intriguingly, TRPV1 expression levels significantly increased in T cells treated with FK506 or B16F10-CS (26.9 ± 1.11 or 23.97 ± 1.04) as compared to resting T cells (16.5 ± 0.52). Moreover, the TRPV1 expression further increased in FK506 or B16F10-CS treated T cells with ConA or TCR stimulation. FK506 or B16F10-CS treated T cells stimulated with ConA or TCR, the TRPV1 expression was 34.53 ± 0.86 or 33.77 ± 0.85 respectively as compared to control ConA activated T cells (28.83 ± 1.19). Similarly, in TCR activated FK506 or B16F10-CS treated T cells, the TRPV1 expression was 33.97 ± 0.77 or 36.1 ± 1.31 respectively as compared to control TCR activated T cells (30.07 ± 0.20) (Figure 1, E). Interestingly, the current findings indicate that the TRPV1 expression also increases significantly in FK506 or B16F10-CS treated T cells as compared to resting T cells, which were further elevated in FK506 or B16F10-CS treated T cells and later stimulated with either ConA or TCR as compared to ConA or TCR activation alone.

### 3.2 Modulation of cytokine response in immunosuppressed T cells

ConA or TCR mediated activation is known to induce robust pro-inflammatory cytokines production in T cells (Majhi et al., 2015; Sahoo et al., 2019, 2018). Conversely, immunosuppressed (FK506 or B16F10-CS treated) T cells stimulated by ConA or TCR show attenuation of pro-inflammatory cytokine response (Almawi et al., 2001; Chen et al., 2019; Sun et al., 2015). Hence, cell culture supernatant were collected at 36 hours to assess the release of cytokines IFN-γ, IL-2 and TNF as studied elsewhere (Majhi et al., 2015; Sahoo et al., 2019, 2018).

For IFN-γ, upon treatment with immunosuppressive FK506 or B16F10-CS T cells, stimulated with ConA, the IFN-Y levels decreased to 486 ± 91.31 pg/ml and 657.5 ± 130.8 pg/ml respectively as compared to control ConA activated cells (6265 ± 752 pg/ml). Similarly, upon treatment with immunosuppressive FK506 or B16F10-CS, T cells stimulated with TCR, the IFN-γ levels decreased to 388.3 ± 111 pg/ml and 704.8 ± 106.7 pg/ml as compared to control TCR activated cells (5472 ± 827 pg/ml) (Supplementary Figure 2 A). These results indicate that both FK506 and B16F10-CS stimulated with Con A or TCR show decreased production of T cell effector cytokine IFN-γ as compared to ConA or TCR activated T cells.

For IL-2, upon treatment with immunosuppressive FK506 or B16F10-CS T cells stimulated with ConA, the IL-2 levels decreased to 181.5 ± 44.29 pg/ml and 295.3 ± 58.97 pg/ml respectively as compared to control ConA activated cells (14081 ± 1740 pg/ml). Similarly, upon treatment with FK506 or B16F10-CS, T cells activated with TCR, the IL-2 levels decreased to 140.8 ± 13.38 pg/ml and 318.8 ± 72.85 pg/ml as compared to control TCR activated cells (12199 ± 565.7 pg/ml) (Supplementary Figure 2, B). These results indicate that both FK506 and B16F10-CS stimulated with Con A or TCR show decreased production of T cell activation cytokine IL-2 as compared to ConA or TCR activated T cells.

For TNF, upon treatment with immunosuppressive FK506 or B16F10-CS, T cells activated with Con A, the TNF levels decreased to 609.3 ± 76.57 pg/ml and 590 ± 55.5 pg/ml respectively as compared to control ConA activated cells (1230 ± 57.36 pg/ml). Similarly, upon treatment with immunosuppressive FK506/B16F10-CS, T cells activated with TCR, the TNF levels decreased to 608.5 ± 35.71 pg/ml and 576.8 ± 43.57 pg/ml as compared to control TCR activated cells (1380 ± 37.38 pg/ml) (Supplementary Figure 2, C). These results indicate that both FK506 and B16F10-CS stimulated with Con A or TCR show decreased production of pro-inflammatory cytokine TNF as compared to ConA or TCR activated T cells. Together these results suggest that the pro-inflammatory cytokines decreased in ConA or TCR stimulated T cells treated with FK506 or B16F10-CS.

### 3.3 Modulation of intracellular Ca^2+^ via TRPV1 channel

5 -IRTX is a potent and specific functional inhibitor of TRPV1 channel and can block TRPV1 directed Ca^2+^ influx (Kumar et al., 2020; Majhi et al., 2015). Here, the intracellular Ca^2+^ levels were assessed by using the Ca^2+^ sensitive dye Fluo-4 AM (Majhi et al., 2015; Sahoo et al., 2019). Accordingly, T cells were pre-treated with 5 -IRTX to determine any change in accumulated intracellular Ca^2+^ during various experimental conditions. Pre-treatment of T cells with TRPV1 functional inhibitor, 5 -IRTX led to reduced calcium influx and subsequently lowers intracellular Ca^2+^ in T cells compared to their corresponding controls. In T cells, treated with FK506, the intracellular Ca^2+^ levels increased markedly compared to resting T cells. Similarly, in T cells, treated with FK506 and stimulated with either ConA or TCR, the Ca^2+^ levels also increased. Further, pre-treatment with 5 -IRTX, leads to marked decrease in accumulated Ca^2+^ levels in T cells (Figure 2). Moreover, in T cells, treated with B16F10-CS, the Ca^2+^ levels also increased compared to resting T cells. Further, pre-treatment with 5 -IRTX, led to decrease in accumulated Ca^2+^ levels in T cells. Further, in T cells, treated with B16F10-CS and stimulated with either ConA or TCR, the Ca^2+^ levels modestly elevated as compared to its corresponding ConA or TCR controls. Further, pre-treatment with 5 -IRTX, led to marked decrease in accumulated Ca^2+^ levels in T cells (Figure 2). As a positive control, the maximum increase in intracellular Ca^2+^ levels were depicted with ionomycin (positive control) (Figure 2, inset). These results highlight that the TRPV1 channel might be acting as a major contributor in elevating intracellular Ca^2+^ levels during both immune-activation and immunosuppression.

**FIGURE 2:**
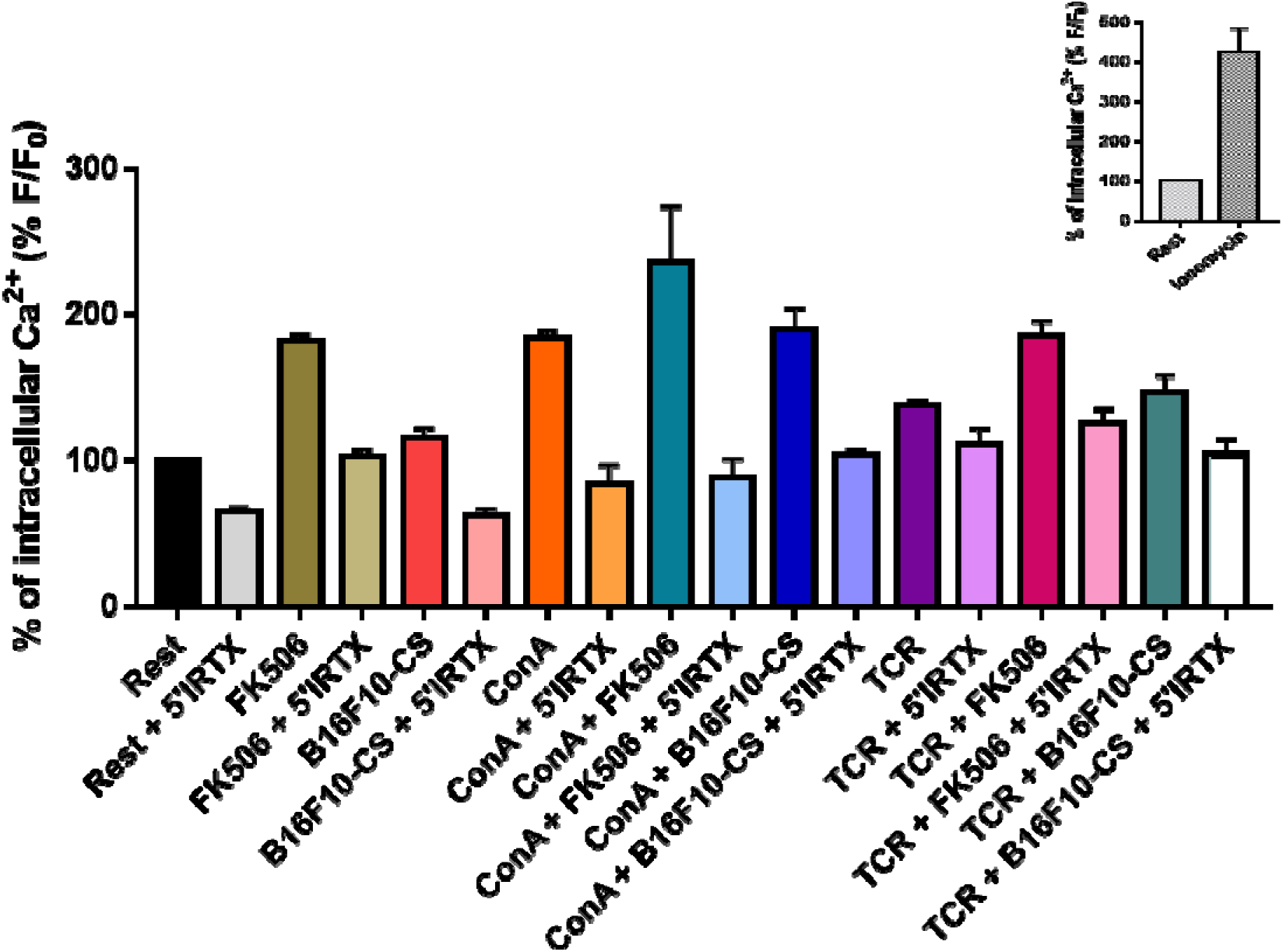
Modulation of intracellular Ca^2+^ via TRPV1 channel. T cells were incubated with calcium sensitive dye, Fluo-4 AM as mentioned in material and methods and acquired via Flow cytometry (FC). Fluo-4 intensity representing intracellular Ca^2+^ has been expressed as percentage normalized to resting control. The various experimental conditions were pre-treated with 5 IRTX, followed by treatment as per experimental conditions. The inset depicts the rise in intracellular Ca^2+^ levels in ionomycin treated T cells. Representative data is of three independent experiments.

### 3.4 Modulation of TRPV1 expression and intracellular Ca^2+^ levels in B16F10-tumor bearing mice, *in vivo*

B16F10, a mouse melanoma cell line, has been injected subcutaneously as described in material and methods. In order to ascertain the modulation of TRPV1 expression and consequent rise in intracellular Ca^2+^ levels in splenic T cells isolated from B16F10-tumor bearing mice, we performed Flow cytometry and calcium influx studies. It has been observed that TRPV1 expression levels significantly increased in T cells isolated from B16F10-tumor bearing mice as compared to control mice (Figure 3, A). Next, we assessed whether the increase in TRPV1 expression levels has also led to the concurrent rise in intracellular Ca^2+^ levels. We found that the basal intracellular Ca^2+^ levels also increased in T cells isolated from B16F10-tumor bearing mice as compared to control mice (Figure 3, B). These results indicate that TRPV1 expression increases in T cells isolated from B16F10-tumor bearing mice and consequently the basal intracellular Ca^2+^ levels were also increased as compared to control mice, *in vivo*.

**FIGURE 3:**
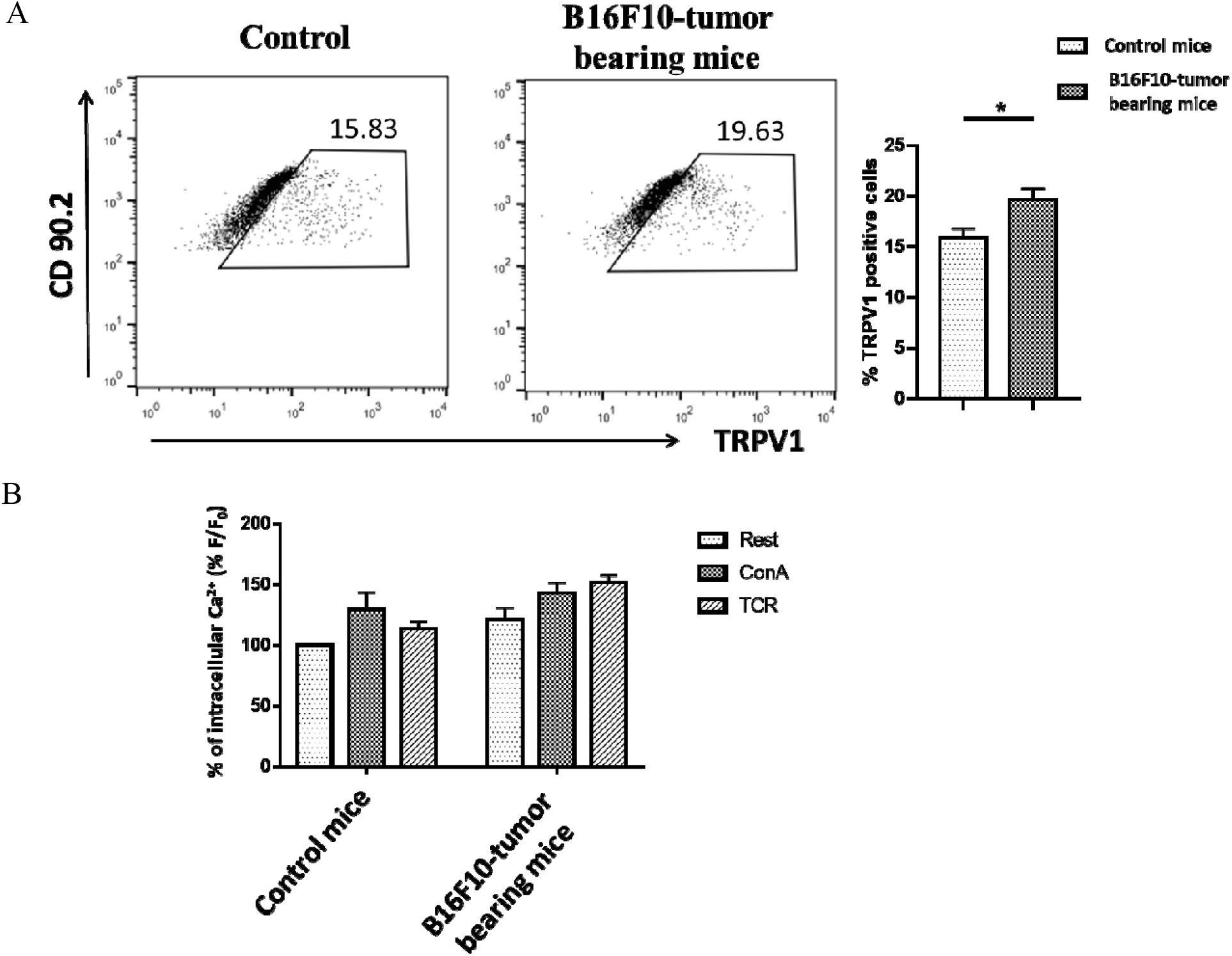
Modulation of TRPV1 expression and intracellular Ca^2+^ levels in B16F10-tumor bearing mice, *in vivo*. Splenic T cells were isolated from tumor bearing mice and analyzed via FC. (A) FC dot-plot depicting TRPV1 expression on T cells along with its corresponding bar diagram. (B) Basal intracellular Ca^2+^ levels in B16F10-tumor bearing mice as compared to control mice. Fluo-4 intensity representing intracellular Ca^2+^ has been expressed as percentage normalized to resting T cells of control mice. Representative data of three independent experiments are shown. *P* < 0.05 was considered as statistically significant difference between the groups (ns, non-significant; * *p* < 0.05; ** *p* < 0.01; *** *p* < 0.001).

## 4. Discussion

TRPV1 is a member of vanilloid group of TRP family and has been attributed in functional expression in a range of immune cells including T cells, macrophages, dendritic cells and NK cells (Assas et al., 2016; Kim et al., 2014; Majhi et al., 2015; Omari et al., 2017). The contributing role of TRPV1 towards Ca^2+^ influx has been very well accredited in T cell development and activation. Moreover, a number of immunosuppressive agents, such as rapamycin, tacrolimus and cyclosporin has also been reported to induce intracellular rise in Ca^2+^ (Bielefeldt et al., 1997; Bultynck et al., 2000; Cameron, Steiner, Roskams, et al., 1995; Cameron, Steiner, Sabatini, et al., 1995; Kanoh et al., 1999; D. MacMillan & McCarron, 2009; Van Acker et al., 2004). However, the role of TRPV1 in calcium influx induced by immunosuppressive agents is not well studied. The current study provides the first evidence that the TRPV1 may also contribute towards FK506 or B16F10-CS driven rise in intracellular calcium associated to immunosuppression of T cells. This study further highlights the modulation of T cell activation, pro-inflammatory cytokine responses and modulation of intracellular Ca^2+^ levels.

TRPV1 is functionally expressed on T cells. TRPV1 has been reported to be upregulated during T cell activation and its associated immune responses (Bertin *et al*., 2014; Majhi *et al*., 2015). Moreover, TRPV1 has also been reported to be actively involved in various inflammatory conditions and pathophysiology (Bujak et al., 2019; Kumar et al., 2020; Majhi et al., 2015; Omari et al., 2017). Upon T cell activation via ConA or TCR, CD69 and CD25 increased significantly. As reported earlier, TRPV1 expression levels increased in activated T cells. Similarly, upon treatment with immunosuppressive FK506 or B16F10-CS, CD69 and CD25 decreased significantly. Unprecedentedly, TRPV1 expression has further increased significantly in immunosuppressed T cells as compared to resting T cells. Furthermore, TRPV1 expression has also been found to be significantly upregulated in FK506 or B16F10-CS treated immunosuppressed T cells stimulated with either ConA or TCR as compared to activated T cells. Moreover, B16F10 tumor-bearing mice also displayed a significant increase in TRPV1 expression levels in T cells. This in general may suggest that during immunosuppressive environment, TRPV1 expression is upregulated in T cells during both *in vitro* and *in vivo*.

T cell activation and suppression are associated with robust cytokine response (Almawi et al., 2001; Majhi et al., 2015; Sahoo et al., 2019, 2018; Sakuma et al., 2000). Upon T cell activation, pro-inflammatory cytokines like IFN-γ, IL-2 and TNF get upregulated. However, T cells treated with immunosuppressive FK506 or B16F10-CS, and stimulated in presence of ConA or TCR, the pro-inflammatory cytokine responses are downregulated. Similar decreasing trend in cytokine response was also reported by others as well (Chen et al., 2019; Sun et al., 2015).

Intracellular Ca^2+^ plays a pivotal role in cell signaling. Modulation of intracellular Ca^2+^ levels is marked with various cellular responses including activation, migration, differentiation and suppression (Aghdasi et al., 2001; Cameron et al., 1995; Feske, 2007; Komada et al., 1996; MacMillan, 2013; Oh-hora & Rao, 2008; Pang et al., 2012; Vig & Kinet, 2009; Wille et al., 2013). T cell activation is marked with rise in intracellular calcium and so as immunosuppression as well. Hence, modulation of TRPV1 channel via 5 -IRTX has been used to validate the contribution of TRPV1 channel towards the FK506 or B16F10-CS driven accumulated intracellular Ca^2+^ levels in T cells. 5 -IRTX is widely used as a functional inhibitor of TRPV1 channel and acts as a functional blocker of TRPV1-mediated Ca^2+^ influx (Kumar et al., 2020; Majhi et al., 2015). In FK506 treated T cells, stimulated with either ConA or TCR, the accumulated intracellular Ca^2+^ levels markedly increased and pre-treatment with 5 -IRTX led to marked decrease in accumulated intracellular Ca^2+^ levels. In B16F10-CS treated T cells, a modest increase in accumulated intracellular Ca^2+^ levels were observed as compared to resting T cells. Further, in other experimental conditions such as, B16F10-CS treated T cells, both in presence or absence of ConA or TCR stimulation, showed a modest increase in accumulated intracellular Ca^2+^ levels as compared to ConA or TCR control. This might be due to the various secreted soluble factors in B16F10-CS released which may regulate increased Ca^2+^ levels (Sun et al., 2015, 2013, 2011; Yang & Carbone, 2004; Zou, 2005). Moreover, B16F10-tumor bearing mice also displayed a significant increase in basal calcium levels in T cells. This in general may suggest that during immunosuppressive environment, the basal calcium levels is upregulated in T cells during both *in vitro* and *in vivo*. These findings indicate that the TRPV1 channel might be acting as a major contributor in elevating intracellular Ca^2+^ levels in T cells. Mechanistically, Ca^2+^ has been reported to activate CaMKK2 which further induces an immunosuppressive microenvironment. Moreover, deletion of CaMKK2 has been reported to attenuate tumor growth, *in vivo* (Racioppi et al., 2019). Further studies are warranted for detailed mechanisms underlying the possible association between elevated calcium and immunosuppressive microenvironment.

In brief, the present study provides evidence that the TRPV1 channel expression is induced on T cells during both immune-activation and immunosuppression. Interestingly, it was found that during immunosuppression, the TRPV1 channel expression and intracellular Ca^2+^ levels in T cells are further elevated. Moreover, the heightened elevation of intracellular Ca^2+^ during experimental immunosuppression was found to be regulated by TRPV1 channel. The current understanding of TRPV1 in T cell activation or suppression is schematically summarized as a proposed working model in figure 4. These findings might also have broad implication to understand the role of TRPV1 channel in various immunosuppressive diseases as well.

**Figure 4:**
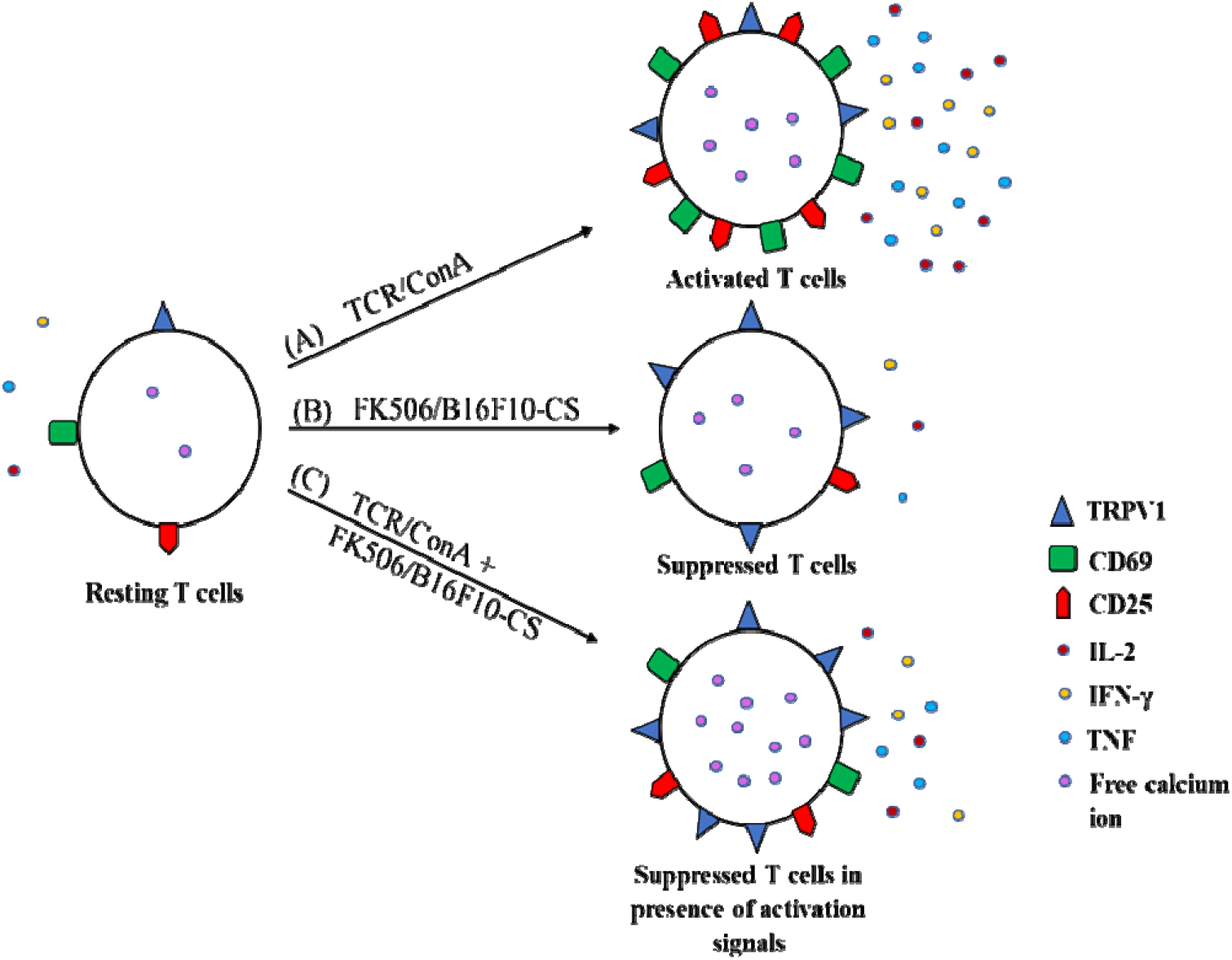
Model depicting functional expression of TRPV1 and intracellular calcium levels in activated and immuno-suppressed T cells. Resting T cells maintain low levels of cytosolic Ca^2+^ and basal levels of TRPV1 expression. (A) Upon activation with either ConA or TCR, both cytosolic Ca^2+^ and TRPV1 are upregulated significantly along with robust effector cytokine responses. (B) Upon treatment with immunosuppressive FK506 or B16F10-CS, both cytosolic Ca^2+^ and TRPV1 expression increases significantly. (C) T cells during experimental immunosuppression (treated with FK506/B16-CS) and stimulated either by ConA or TCR, although shows decreased cytokine response, may show significantly higher cytosolic Ca^2+^ and TRPV1 expression.

## Acknowledgement

We are thankful to Dr. Chandan Goswami, SBS/NISER, Bhubaneswar, India, for the necessary advices for the work. We are also thankful to the Animal House Facility and Flow Cytometry Facility of NISER for their support.

## Author Contributions

Conceptualization: P Sanjai Kumar, Tathagata Mukherjee, Subhasis Chattopadhyay; Methodology: P Sanjai Kumar, Tathagata Mukherjee, Somlata Khamaru, Dalai Jupiter Nanda Kishore, Subhransu Sekhar Sahoo; Formal analysis and investigation: P Sanjai Kumar, Tathagata Mukherjee, Subhasis Chattopadhyay; Writing - original draft preparation: P Sanjai Kumar, Tathagata Mukherjee, Subhasis Chattopadhyay; Writing - review and editing: P Sanjai Kumar, Tathagata Mukherjee, Somlata Khamaru, Dalai Jupiter Nanda Kishore, Subhransu S. Sahoo, Subhasis Chattopadhyay; Funding acquisition: Subhasis Chattopadhyay; Resources: Subhasis Chattopadhyay; Supervision: Subhasis Chattopadhyay

## References

Acharya, T. K., Tiwari, A., Majhi, R. K., & Goswami, C. (2020). TRPM8 channel augments T cell activation and proliferation. Cell Biology International, (November). https://doi.org/10.1002/cbin.11483

Aghdasi, B., Ye, K., Resnick, A., Huang, A., Ha, H. C., Guo, X., … Snyder, S. H. (2001). Physiologic Regulator of the Cell Cycle. Proceedings of the National Academy of Science, 98(5), 2425–2430.

Almawi, W. Y., Assi, J. W., Chudzik, D. M., Jaoude, M. M. A., & Rieder, M. J. (2001). Inhibition of cytokine production and cytokine-stimulated T-cell activation by FK506 (Tacrolimus)1. Cell Transplantation, 10(7), 615–623. https://doi.org/10.3727/000000001783986387

Amantini, C., Farfariello, V., Cardinali, C., Morelli, M. B., Marinelli, O., Nabissi, M., … Santoni, G. (2017). The TRPV1 ion channel regulates thymocyte differentiation by modulating autophagy and proteasome activity. Oncotarget, 8(53), 90766–90780. https://doi.org/10.18632/oncotarget.21798

Ando, Y., Yasuoka, C., Mishima, T., Ikematsu, T., Uede, T., Matsunaga, T., & Inobe, M. (2014). Concanavalin A-mediated T cell proliferation is regulated by herpes virus entry mediator costimulatory molecule. In Vitro Cellular and Developmental Biology - Animal, 50(4), 313–320. https://doi.org/10.1007/s11626-013-9705-2

Annett, S., Moore, G., & Robson, T. (2020). FK506 binding proteins and inflammation related signalling pathways; basic biology, current status and future prospects for pharmacological intervention. Pharmacology & Therapeutics, 215, 107623. https://doi.org/10.1016/j.pharmthera.2020.107623

Assas, B. M., Wakid, M. H., Zakai, H. A., Miyan, J. A., & Pennock, J. L. (2016). Transient receptor potential vanilloid 1 expression and function in splenic dendritic cells: A potential role in immune homeostasis. Immunology, 147(3), 292–304. https://doi.org/10.1111/imm.12562

Bassi, M. S., Gentile, A., Iezzi, E., Zagaglia, S., Musella, A., Simonelli, I., … Buttari, F. (2019). Transient receptor potential vanilloid 1 modulates central inflammation in multiple sclerosis. Frontiers in Neurology, 10(JAN), 1–8. https://doi.org/10.3389/fneur.2019.00030

Bertin, S., Aoki-Nonaka, Y., De Jong, P. R., Nohara, L. L., Xu, H., Stanwood, S. R., … Raz, E. (2014). The ion channel TRPV1 regulates the activation and pro-inflammatory properties of CD4+T cells. Nature Immunology, 15(11), 1055–1063. https://doi.org/10.1038/ni.3009

Bielefeldt, K., Sharma, R. V., Whiteis, C., Yedidag, E., & Abboud, F. M. (1997). Tacrolimus (FK506) modulates calcium release and contractility of intestinal smooth muscle. Cell Calcium, 22(6), 507–514. https://doi.org/10.1016/S0143-4160(97)90078-6

Birx, D. L., Berger, M., & Fleisher, T. A. (1984). The interference of T cell activation by calcium channel blocking agents. The Journal of Immunology, 133(6).

Bujak, J. K., Kosmala, D., Szopa, I. M., Majchrzak, K., & Bednarczyk, P. (2019, October 16). Inflammation, Cancer and Immunity—Implication of TRPV1 Channel. Frontiers in Oncology, Vol. 9, p. 1087. Frontiers Media S.A. https://doi.org/10.3389/fonc.2019.01087

Bultynck, G., De Smet, P., Weidema, A. F., Ver Heyen, M., Maes, K., Callewaert, G., … De Smedt, H. (2000). Effects of the immunosuppressant FK506 on intracellular Ca2+ release and Ca2+ accumulation mechanisms. Journal of Physiology, 525(3), 681–693. https://doi.org/10.1111/j.1469-7793.2000.t01-1-00681.x

Burghoff, S., Gong, X., Viethen, C., Jacoby, C., Flögel, U., Bongardt, S., … Schrader, J. (2014). Growth and metastasis of B16-F10 melanoma cells is not critically dependent on host CD73 expression in mice. BMC Cancer, 14(1), 898. https://doi.org/10.1186/1471-2407-14-898

Cameron, A. M., Steiner, J. P., Roskams, A. J., Ali, S. M., Ronnettt, G. V., & Snyder, S. H. (1995). Calcineurin associated with the inositol 1,4,5-trisphosphate receptor-FKBP12 complex modulates Ca2+ flux. Cell, 83(3), 463–472. https://doi.org/10.1016/0092-8674(95)90124-8

Cameron, A. M., Steiner, J. P., Sabatini, D. M., Kaplin, A. I., Walensky, L. D., & Snyder, S. H. (1995). Immunophilin FK506 binding protein associated with inositol 1,4,5-trisphosphate receptor modulates calcium flux. Proceedings of the National Academy of Sciences of the United States of America, 92(5), 1784–1788. https://doi.org/10.1073/pnas.92.5.1784

Chen, Y.-Q., Li, P.-C., Pan, N., Gao, R., Wen, Z.-F., Zhang, T.-Y., … Wang, L.-X. (2019). c. Journal for ImmunoTherapy of Cancer, 7(1), 178. https://doi.org/10.1186/s40425-019-0646-5

Feske, S. (2007). Calcium signalling in lymphocyte activation and disease. Nature Reviews Immunology, 7(9), 690–702. https://doi.org/10.1038/nri2152

Gao, R., Ma, J., Wen, Z., Yang, P., Zhao, J., Xue, M., … Pan, N. (2018). Tumor cell-released autophagosomes (TRAP) enhance apoptosis and immunosuppressive functions of neutrophils. OncoImmunology, 7(6), 1–11. https://doi.org/10.1080/2162402X.2018.1438108

Gees, M., Colsoul, B., & Nilius, B. (2010). The role of transient receptor potential cation channels in Ca2+ signaling. Cold Spring Harbor Perspectives in Biology, 2(10), a003962–a003962. https://doi.org/10.1101/cshperspect.a003962

Kanoh, S., Kondo, M., Tamaoki, J., Shirakawa, H., Aoshiba, K., Miyazaki, S., … Nagai, A. (1999). Effect of FK506 on ATP-induced intracellular calcium oscillations in cow tracheal epithelium. American Journal of Physiology - Lung Cellular and Molecular Physiology, 276(6 20-6), 891–899. https://doi.org/10.1152/ajplung.1999.276.6.l891

Kim, H. S., Kwon, H. J., Kim, G. E., Cho, M. H., Yoon, S. Y., Davies, A. J., … Kim, Y. K. (2014). Attenuation of natural killer cell functions by capsaicin through a direct and TRPV1-independent mechanism: Capsaicin-induced NK cell dysfunction. Carcinogenesis, 35(7), 1652–1660. https://doi.org/10.1093/carcin/bgu091

Kita, T., Uchida, K., Kato, K., Suzuki, Y., Tominaga, M., & Yamazaki, J. (2019). FK506 (tacrolimus) causes pain sensation through the activation of transient receptor potential ankyrin 1 (TRPA1) channels. The Journal of Physiological Sciences, 69(2), 305–316. https://doi.org/10.1007/s12576-018-0647-z

Komada, H., Nakabayashi, H., Hara, M., & Izutsu, K. (1996). Early calcium signaling and calcium requirements for the IL-2 receptor expression and IL-2 production in stimulated lymphocytes. Cellular Immunology, 173(2), 215–220. https://doi.org/10.1006/cimm.1996.0270

Kumar, P. S., Nayak, T. K., Mahish, C., Sahoo, S. S., & Radhakrishnan, A. (2020). Inhibition of transient receptor potential vanilloid 1 (TRPV1) channel regulates chikungunya virus infection in macrophages. Archives of Virology, 1(0123456789), 17–19. https://doi.org/10.1007/s00705-020-04852-8

Kume, H., & Tsukimoto, M. (2019). TRPM8 channel inhibitor AMTB suppresses murine T-cell activation induced by T-cell receptor stimulation, concanavalin A, or external antigen restimulation. Biochemical and Biophysical Research Communications, 509(4), 918–924. https://doi.org/10.1016/j.bbrc.2019.01.004

Kusmartsev, S., & Gabrilovich, D. I. (2002, August 1). Immature myeloid cells and cancer-associated immune suppression. Cancer Immunology, Immunotherapy, Vol. 51, pp. 293–298. https://doi.org/10.1007/s00262-002-0280-8

Kusmartsev, S., Nefedova, Y., Yoder, D., & Gabrilovich, D. I. (2004). Antigen-Specific Inhibition of CD8 + T Cell Response by Immature Myeloid Cells in Cancer Is Mediated by Reactive Oxygen Species. The Journal of Immunology, 172(2), 989–999. https://doi.org/10.4049/jimmunol.172.2.989

Langer, F., Amirkhosravi, A., Ingersoll, S. B., Walker, J. M., Spath, B., Eifrig, B., … Francis, J. L. (2006). Experimental metastasis and primary tumor growth in mice with hemophilia A. Journal of Thrombosis and Haemostasis : JTH, 4(5), 1056–1062. https://doi.org/10.1111/j.1538-7836.2006.01883.x

Mackler, B. F., Wolstencroft, R. A., & Dumonde, D. C. (1972). Concanavalin a as an inducer of human lymphocyte mitogenic factor. Nature New Biology, 239(92), 139–142. https://doi.org/10.1038/newbio239139a0

MacMillan, D., & McCarron, J. G. J. (2009). Regulation by FK506 and rapamycin of Ca2+ release from the sarcoplasmic reticulum in vascular smooth muscle: the role of FK506 binding proteins and mTOR. British Journal of Pharmacology, 158(4), 1112–1120. https://doi.org/10.1111/j.1476-5381.2009.00369.x

MacMillan, Debbi. (2013). FK506 binding proteins: Cellular regulators of intracellular Ca 2+ signalling. European Journal of Pharmacology, 700(1–3), 181–193. https://doi.org/10.1016/j.ejphar.2012.12.029

Majhi, R. K., Sahoo, S. S., Yadav, M., Pratheek, B. M., Chattopadhyay, S., & Goswami, C. (2015). Functional expression of TRPV channels in T cells and their implications in immune regulation. FEBS Journal, 282(14), 2661–2681. https://doi.org/10.1111/febs.13306

Nakabayashi, H., Komada, H., Yoshida, T., Takanari, H., & Izutsu, K. (1992). Lymphocyte calmodulin and its participation in the stimulation of T lymphocytes by mitogenic lectins. Biology of the Cell, 75(1), 55–59. https://doi.org/10.1016/0248-4900(92)90124-J

Nayak, T. K., Mamidi, P., Kumar, A., Singh, L. P. K., Sahoo, S. S., Chattopadhyay, S., & Chattopadhyay, S. (2017). Regulation of viral replication, apoptosis and pro-inflammatory responses by 17-aag during chikungunya virus infection in macrophages. Viruses, 9(1). https://doi.org/10.3390/v9010003

Nilius, B. (2007). TRP channels in disease. Biochimica et Biophysica Acta - Molecular Basis of Disease, 1772(8), 805–812. https://doi.org/10.1016/j.bbadis.2007.02.002

Oh-hora, M., & Rao, A. (2008, June 1). Calcium signaling in lymphocytes. Current Opinion in Immunology, Vol. 20, pp. 250–258. Elsevier Current Trends. https://doi.org/10.1016/j.coi.2008.04.004

Omar, S., Clarke, R., Abdullah, H., Brady, C., Corry, J., Winter, H., … Cosby, S. L. (2017). Respiratory virus infection upregulates TRPV1, TRPA1 and ASICS3 receptors on airway cells. PLoS ONE, 12(2), 1–21. https://doi.org/10.1371/journal.pone.0171681

Omari, S. A., Adams, M. J., & Geraghty, D. P. (2017). TRPV1 Channels in Immune Cells and Hematological Malignancies. In Advances in Pharmacology (1st ed., Vol. 79, pp. 173–198). Elsevier Inc. https://doi.org/10.1016/bs.apha.2017.01.002

Overwijk, W. W., & Restifo, N. P. (2001). B16 as a Mouse Model for Human Melanoma. Current Protocols in Immunology, 39(1), Unit 20.1. https://doi.org/10.1002/0471142735.im2001s39

Pang, B., Shin, D. H., Park, K. S., Huh, Y. J., Woo, J., Zhang, Y. H., … Kim, S. J. (2012). Differential pathways for calcium influx activated by concanavalin A and CD3 stimulation in Jurkat T cells. Pflugers Archiv European Journal of Physiology, 463(2), 309–318. https://doi.org/10.1007/s00424-011-1039-x

Racioppi, L., Nelson, E. R., Huang, W., Mukherjee, D., Lawrence, S. A., Lento, W., … McDonnell, D. P. (2019). CaMKK2 in myeloid cells is a key regulator of the immune-suppressive microenvironment in breast cancer. Nature Communications, 10(1), 1–16. https://doi.org/10.1038/s41467-019-10424-5

Sahoo, S. S., Majhi, R. K., Tiwari, A., Acharya, T., Kumar, P. S., Saha, S., … Chattopadhyay, S. (2019). Transient receptor potential ankyrin1 channel is endogenously expressed in T cells and is involved in immune functions. Bioscience Reports, 39(9), BSR20191437. https://doi.org/10.1042/BSR20191437

Sahoo, S. S., Pratheek, B. M., Meena, V. S., Nayak, T. K., Kumar, P. S., Bandyopadhyay, S., … Chattopadhyay, S. (2018). VIPER regulates naive T cell activation and effector responses: Implication in TLR4 associated acute stage T cell responses. Scientific Reports, 8(1), 1–9. https://doi.org/10.1038/s41598-018-25549-8

Sakuma, S., Kato, Y., Nishigaki, F., Sasakawa, T., Magari, K., Miyata, S., … Goto, T. (2000). FK506 potently inhibits T cell activation induced TNF-α and IL-1β production in vitro by human peripheral blood mononuclear cells. British Journal of Pharmacology, 130(7), 1655–1663. https://doi.org/10.1038/sj.bjp.0703472

Samanta, A., Hughes, T. E. T., & Moiseenkova-Bell, V. Y. (2018). Transient receptor potential (TRP) channels. In Subcellular Biochemistry (Vol. 87, pp. 141–165). Springer New York. https://doi.org/10.1007/978-981-10-7757-9_6

Silva, R. O., Bingana, R. D., Sales, T. M. A. L., Moreira, R. L. R., Costa, D. V. S., Sales, K. M. O., … Souza, M. H. L. P. (2018). Role of TRPV1 receptor in inflammation and impairment of esophageal mucosal integrity in a murine model of nonerosive reflux disease. Neurogastroenterology and Motility, 30(8), 1–8. https://doi.org/10.1111/nmo.13340

Sun, L. X., Li, W. D., Lin, Z. Bin, Duan, X. S., Xing, E. H., Jiang, M. M., … Lu, J. (2015). Cytokine production suppression by culture supernatant of B16F10 cells and amelioration by Ganoderma lucidum polysaccharides in activated lymphocytes. Cell and Tissue Research, 360(2), 379–389. https://doi.org/10.1007/s00441-014-2083-6

Sun, L. X., Lin, Z. Bin, Duan, X. S., Lu, J., Ge, Z. H., Li, M., … Li, W. D. (2013). Ganoderma lucidum polysaccharides counteract inhibition on CD71 and FasL expression by culture supernatant of B16F10 cells upon lymphocyte activation. Experimental and Therapeutic Medicine, 5(4), 1117–1122. https://doi.org/10.3892/etm.2013.931

Sun, L. X., Lin, Z. Bin, Duan, X. S., Lu, J., Ge, Z. H., Li, X. J., … Li, W. D. (2011). Ganoderma lucidum polysaccharides antagonize the suppression on lymphocytes induced by culture supernatants of B16F10 melanoma cells. Journal of Pharmacy and Pharmacology, 63(5), 725–735. https://doi.org/10.1111/j.2042-7158.2011.01266.x

Van Acker, K., Bultynck, G., Rossi, D., Sorrentino, V., Boens, N., Missiaen, L., … Callewaert, G. (2004). The 12 kDa FK506-binding protein, FKBP12, modulates the Ca 2+-flux properties of the type-3 ryanodine receptor. Journal of Cell Science, 117(7), 1129–1137. https://doi.org/10.1242/jcs.00948

Vig, M., & Kinet, J. P. (2009, January 17). Calcium signaling in immune cells. Nature Immunology, Vol. 10, pp. 21–27. https://doi.org/10.1038/ni.f.220

Watman, N. P., Crespo, L., Davis, B., Poiesz, B. J., & Zamkoff, K. W. (1988). Differential effect on fresh and cultured T cells of PHA-induced changes in free cytoplasmic calcium: Relation to IL-2 receptor expression, IL-2 production, and proliferation. Cellular Immunology, 111(1), 158–166. https://doi.org/10.1016/0008-8749(88)90060-3

Wen, Z. F., Liu, H., Gao, R., Zhou, M., Ma, J., Zhang, Y., … Wang, L. X. (2018). Tumor cell-released autophagosomes (TRAPs) promote immunosuppression through induction of M2-like macrophages with increased expression of PD-L1. Journal for ImmunoTherapy of Cancer, 6(1), 1–16. https://doi.org/10.1186/s40425-018-0452-5

Wille, A. C. M., Chaves-Moreira, D., Trevisan-Silva, D., Magnoni, M. G., Boia-Ferreira, M., Gremski, L. H., … Veiga, S. S. (2013). Modulation of membrane phospholipids, the cytosolic calcium influx and cell proliferation following treatment of B16-F10 cells with recombinant phospholipase-D from Loxosceles intermedia (brown spider) venom. Toxicon, 67, 17–30. https://doi.org/10.1016/j.toxicon.2013.01.027

Yang, L., & Carbone, D. P. (2004). Tumor-host immune interactions and dendritic cell dysfunction. Advances in Cancer Research, 92, 13–27. https://doi.org/10.1016/S0065-230X(04)92002-7

Zhou, M., Wen, Z., Cheng, F., Ma, J., Li, W., Ren, H., … Wang, L. (2016). Tumor-released autophagosomes induce IL-10-producing B cells with suppressive activity on T lymphocytes via TLR2-MyD88-NF-κ B signal pathway. OncoImmunology, 5(7), 1–14. https://doi.org/10.1080/2162402X.2016.1180485

Zou, W. (2005, April 18). Immunosuppressive networks in the tumour environment and their therapeutic relevance. Nature Reviews Cancer, Vol. 5, pp. 263–274. https://doi.org/10.1038/nrc1586

