## Supplementary Data for "Elevation of TRPV1 expression on T cells during experimental immunosuppression"

**experimental immunosuppression**

**P Sanjai Kumar^1,#^, Tathagata Mukherjee^1,#^, Somlata Khamaru^1^, Dalai Jupiter Nanda Kishore^1^, Saurabh Chawla^1^, Subhransu Sekhar Sahoo^1^, Subhasis Chattopadhyay^1,*^**

^1^	School of Biological Sciences, National Institute of Science Education & Research, Bhubaneswar, HBNI, Jatni, Khurda, Odisha 752050, India;

\#	Equal contribution

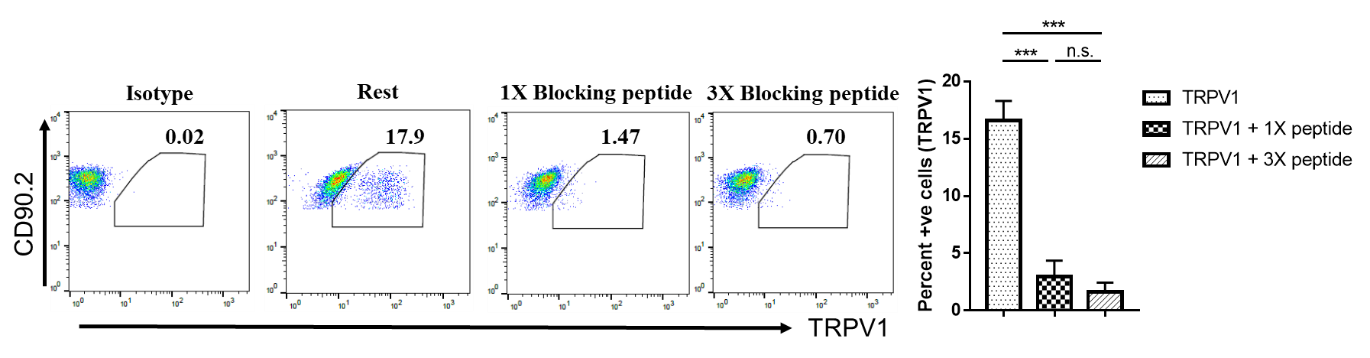

**SUPPLEMENTARY FIGURE 1: TRPV1 expression in purified mouse T cells.**

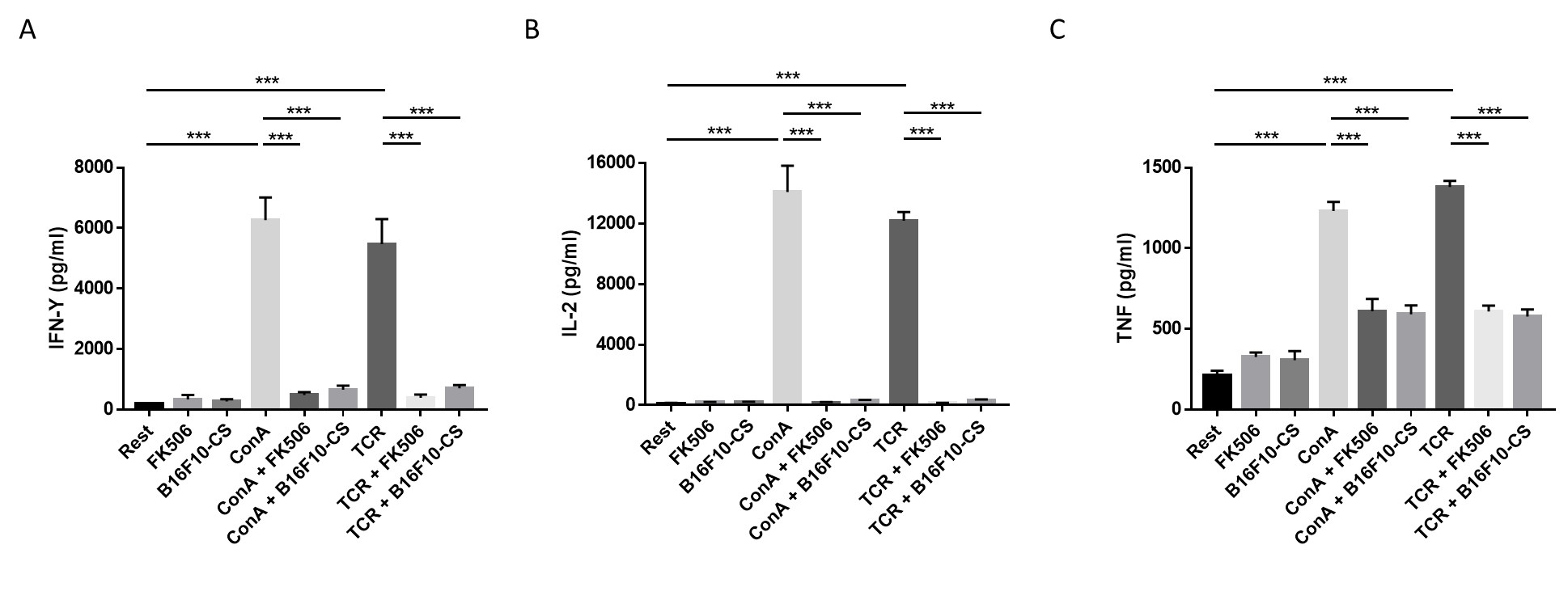
Resting T cells were stained with anti-TRPV1 antibody for 30 mins in FACS buffer followed by secondary antibody staining and acquired via Flow cytometry (FC). Representative dot-plot showing TRPV1 expression in resting T cells in presence or absence of control blocking peptide along with bar diagram. Representative data of three independent experiments are shown. *p* < 0.05 was considered as statistically significant difference between the groups (* *p* < 0.05; * * *p* < 0.01; * * * *p* < 0.001).

**SUPPLEMENTARY FIGURE 2:**  **Modulation of cytokine response in immunosuppressed T cells.**

T cells were treated with FK506 (5µg/mL) or B16F10-CS (20% of final volume) in presence or absence of ConA (5µg/mL) or TCR (2µg/mL of each anti-CD3 and CD28). The supernatant was collected at 36h and sandwich ELISA was performed to quantitate secreted cytokines, such as, (A) IFN-Y, (B) IL-2 and (C) TNF. Representative data of three independent experiments are shown. *p* < 0.05 was considered as statistically significant difference between the groups (*p* < 0.001).
